# *wspA* mutation mediated *Pseudomonas aeruginosa* rugose small colony variant and its hyper-biofilm

**DOI:** 10.1101/2025.09.15.676455

**Authors:** Qing Yu, Yue Li, Peiqi Wang, Chen Zhang, Hanming Ni, Yisha Zhang, Yushan Cui, Mengyu Zhou, Jun Ni, Linyue Zhang, Xiaoting Hua, Yunsong Yu, Qiucheng Shi, Xiaoxing Du

## Abstract

*Pseudomonas aeruginosa* is a major multidrug-resistant pathogen whose biofilm formation complicates treatment. *P. aeruginosa* <u>r</u>ugose <u>s</u>mall <u>c</u>olony <u>v</u>ariants (RSCVs), characterized by enhanced biofilm production and resistance, pose significant clinical challenges. However, the formation mechanisms remain unclear in clinical strains. Paired clinical *P. aeruginosa* isolates AR8023-1 (wild-type) and AR8023-2 (RSCV) were collected from urine samples of a neurosurgery inpatient chronologically. The genome comparison was performed using Breseq, and plasmid complementation was performed to construct complemented strain (AR8023-2^RifR^::*wspA*^AR8023-1^). Then, all the aforementioned strains were examined RSCV phenotype, biofilm formation, and motility, and gene expression differences induced by the *wspA* mutation were analyzed via transcriptome sequencing, followed by quantification of c-di-GMP. In addition, we assessed antimicrobial susceptibility of the strains under both planktonic and biofilm conditions. The results showed that both AR8023-1 and AR8023-2 were identified as ST3420. AR8023-1 exhibited typical smooth morphology, while AR8023-2 displayed a RSCV phenotype. A single SNP difference was identified between the two strains, characterized by a deletion of glutamine at position 289 (CAG triplet deletion) in the *wspA* gene of strain AR8023-2. The complement isolates restored wild-type morphology, motility, and biofilm formation. Transcriptomics revealed a significant upregulation of c-di-GMP metabolic genes in RSCVs (*P* < 0.001), indicating that the WspAΔ289Q mutation activates diguanylate cyclase (DGC) activity, thereby elevating c-di-GMP synthesis. Intracellular c-di-GMP levels were significantly higher in RSCVs than those in WT and complement isolates, respectively (*P* < 0.001). Consequently, biofilm susceptibility testing demonstrated the β-lactams MBIC of RSCVs was ≥8–512-fold higher than those of WT and complement isolates. This study identifies a clinical WspAΔ289Q mutation that elevates c-di-GMP, driving RSCV formation and biofilm-mediated resistance.

**IMPORTANCE:** *P. aeruginosa* is a leading multidrug-resistant pathogen whose biofilm formation capability greatly complicates treatment outcomes. This study identifies a novel clinical mutation in the *wspA* gene (WspAΔ289Q) that drives the emergence of rugose small colony variants (RSCVs), a phenotype linked to heightened biofilm production and antimicrobial resistance. We demonstrate that this mutation upregulates cyclic di-GMP synthesis, leading to increased biofilm formation and markedly reduced susceptibility to β-lactam antibiotics under biofilm conditions. These results provide important insights into the genetic basis of RSCV development in clinical settings and highlight the role of c-di-GMP signaling in biofilm-associated resistance. Our findings underscore the need to target c-di-GMP pathways as a potential strategy for combating persistent *P. aeruginosa* infections.

## INTRODUCTION

*Pseudomonas aeruginosa* poses a severe antimicrobial resistance (AMR) threat (CDC, 2019). Globally, it is a leading bacterial cause of death, with an age-standardized mortality rate of 7.4 per 100,000 ^[1,2]^. In China (CHINET 2024), it accounts for 9.0% of Gram-negative isolates and exhibits high multidrug resistance, notably a 22.0% carbapenem resistance rate, significantly increasing treatment failure risk in critically ill patients. The CDC highlights its capacity to evade host immunity and antibiotics through biofilm formation and phenotypic plasticity, such as the emergence of rugose <u>s</u>mall <u>c</u>olony <u>v</u>ariants (RSCVs) ^[3]^. RSCVs, frequently isolated from chronic infections like cystic fibrosis ^[4]^, are linked to recurrent infections and treatment failure due to enhanced biofilm barriers and metabolic dormancy. However, the molecular mechanisms driving RSCV formation remain incompletely understood, hindering targeted intervention strategies.

The Wsp (Wrinkly Spreader Phenotype) system, a key signaling network regulating surface attachment ^[5]^, utilizes its core transmembrane sensor protein WspA to modulate bacterial adhesion and biofilm formation via c-di-GMP signaling cascades ^[6,7]^. While *wspA* deletion induces phenotypic switching ^[8]^, critical knowledge gaps persist: The specific structural domains and conformational dynamics of WspA in activating downstream c-di-GMP synthesis are unelucidated; Research relies heavily on lab reference strains (e.g., PAO1), lacking validation of mutation functional heterogeneity in clinical isolates; the clinical relevance of WspA mutations, particularly their impact on *in vivo* pathogenicity and resistance profiles, is underexplored, limiting their utility as biomarkers or therapeutic targets.

To address these knowledge gaps, we implemented an integrated multi-omics strategy utilizing paired clinical isolates of *P. aeruginosa* (AR8023-1: wild-type; AR8023-2: RSCV mutant) obtained from a tertiary hospital in Ningbo. This approach encompassed Illumina-Nanopore sequencing, Breseq-based genomic analysis, *wspA* mutant construction, transcriptomic profiling, c-di-GMP quantification, biofilm phenotyping, motility assays, and antimicrobial susceptibility testing.

This study elucidates the role of the WspAΔ289Q mutation in RSCV formation and its impact on antibiotic resistance, thereby identifying biomarkers for surveillance and informing anti-biofilm therapies targeting c-di-GMP signaling.

## RESULTS

### Clinical strain characterization

In 2021, two *P. aeruginosa* strains (designated AR8023-1 and AR8023-2) were isolated from sequential urine culture samples of a postoperative cerebellar infarction patient hospitalized in a tertiary hospital in Ningbo, China. Significantly, significant phenotypic divergences were observed: On blood agar plates, AR8023-1 exhibited large, smooth, moist, and yellowish-green colonies (Fig. 1A), whereas AR8023-2 formed small, dry, wrinkled colonies with a similar pigmentation (Fig. 1B). Under stereomicroscopy, AR8023-1 showed diffuse adherent growth (Fig. 1D), while AR8023-2 developed compact, matrix-encapsulated three-dimensional biofilm structures (Fig. 1E). Notably, the morphological and structural features of AR8023-2 align with the defining characteristics of rugose small colony variants (RSCVs).

**Fig. 1.**
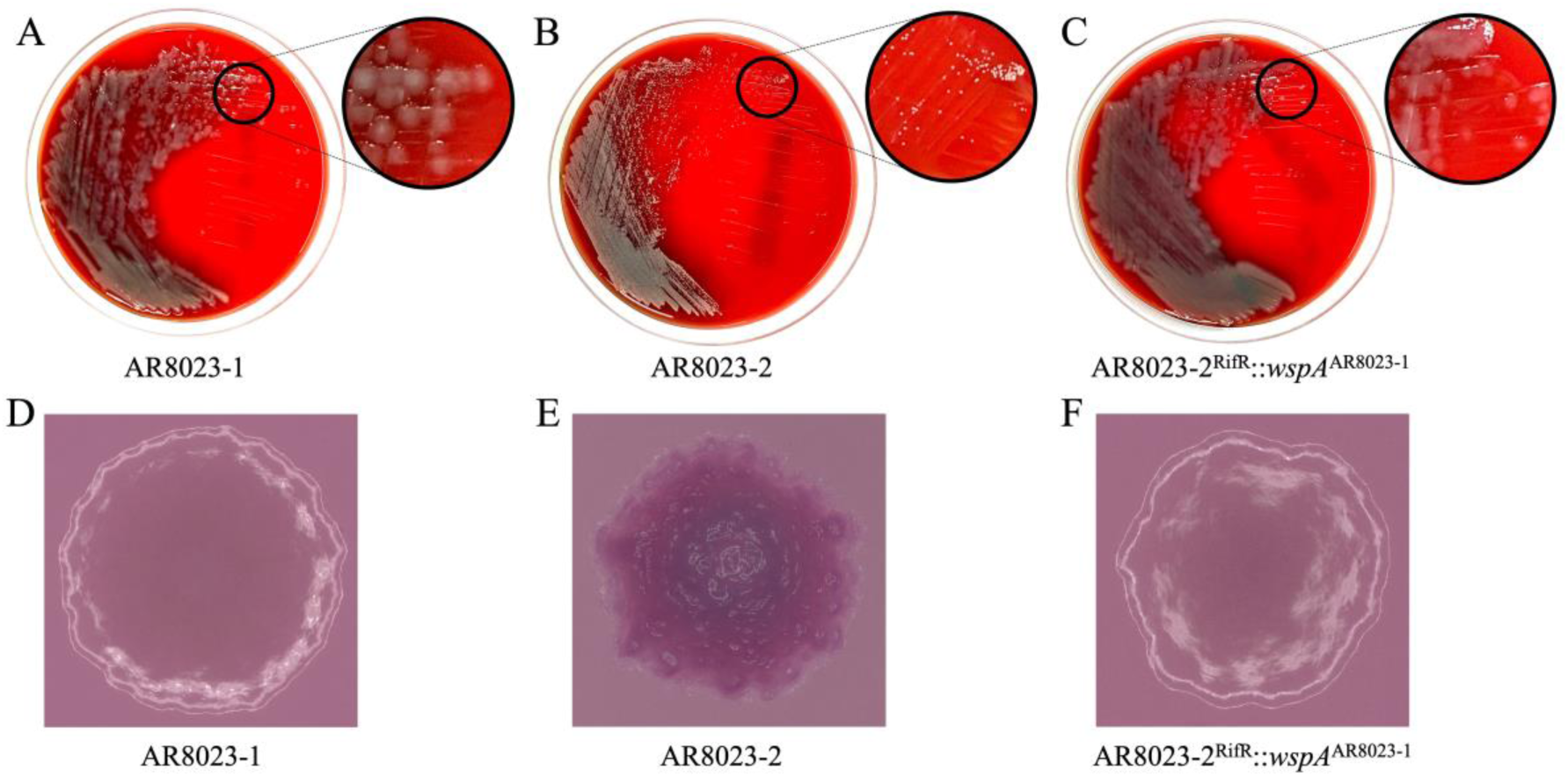
Colony morphology and microscopic features of *P. aeruginosa* strains. (A-C) Colony morphology of *P. aeruginosa* strains cultured overnight at 37 °C on blood agar plates. The large black circle denotes a partial enlarged view of the specified region. (D-F) Colony morphology was assessed by spot-inoculating 1 μL of overnight bacterial culture onto modified LB agar plates, followed by 24-hour incubation at 37°C. Images were acquired using a KEYENCE VHX-7000 digital microscope at 24-hour post-inoculation.

### Genomic identification of WspAΔ289Q mutation and phenotypic validation

Whole-genome sequencing revealed that the clinical isolates AR8023-1 and AR8023-2 (both identified as ST3420) differed by only a single nucleotide polymorphism (SNP), indicating extremely close phylogenetic relatedness and significant homology similarity between the two strains ^[9]^. Compared to AR8023-1, AR8023-2 harbored a CAG trinucleotide repeat deletion at positions 865–867 of the *wspA* gene, resulting in the loss of a glutamine residue at position 289 (ΔQ289) in the encoded protein.

In this study, AR8023-1 and AR8023-2 were designated as the wild-type and mutant strain, respectively. Plasmid-based complementation of AR8023-2 (complemented strain) with *wspA* from AR8023-1 (pGK_WspA^8023-1^) restored wild-type colony morphology (Fig. 1C), confirming that the RSCV phenotype of AR8023-2 directly results from the Δ289Q mutation in WspA. This unequivocally establishes WspA structural integrity as the key regulator of colony morphology in *P. aeruginosa*.

### Biofilm and Motility defect

Firstly, biofilm biomass was measured for all strains (Fig. 2D). Biofilm biomass was significantly greater in AR8023-2 (3.90 ± 0.04) compared to AR8023-1 (0.80 ± 0.05, *P* < 0.001) and AR8023-2^RifR^::*wspA*^AR8023-1^ (1.01 ± 0.05, *P* < 0.001) (Fig. 2E).

**Fig. 2.**
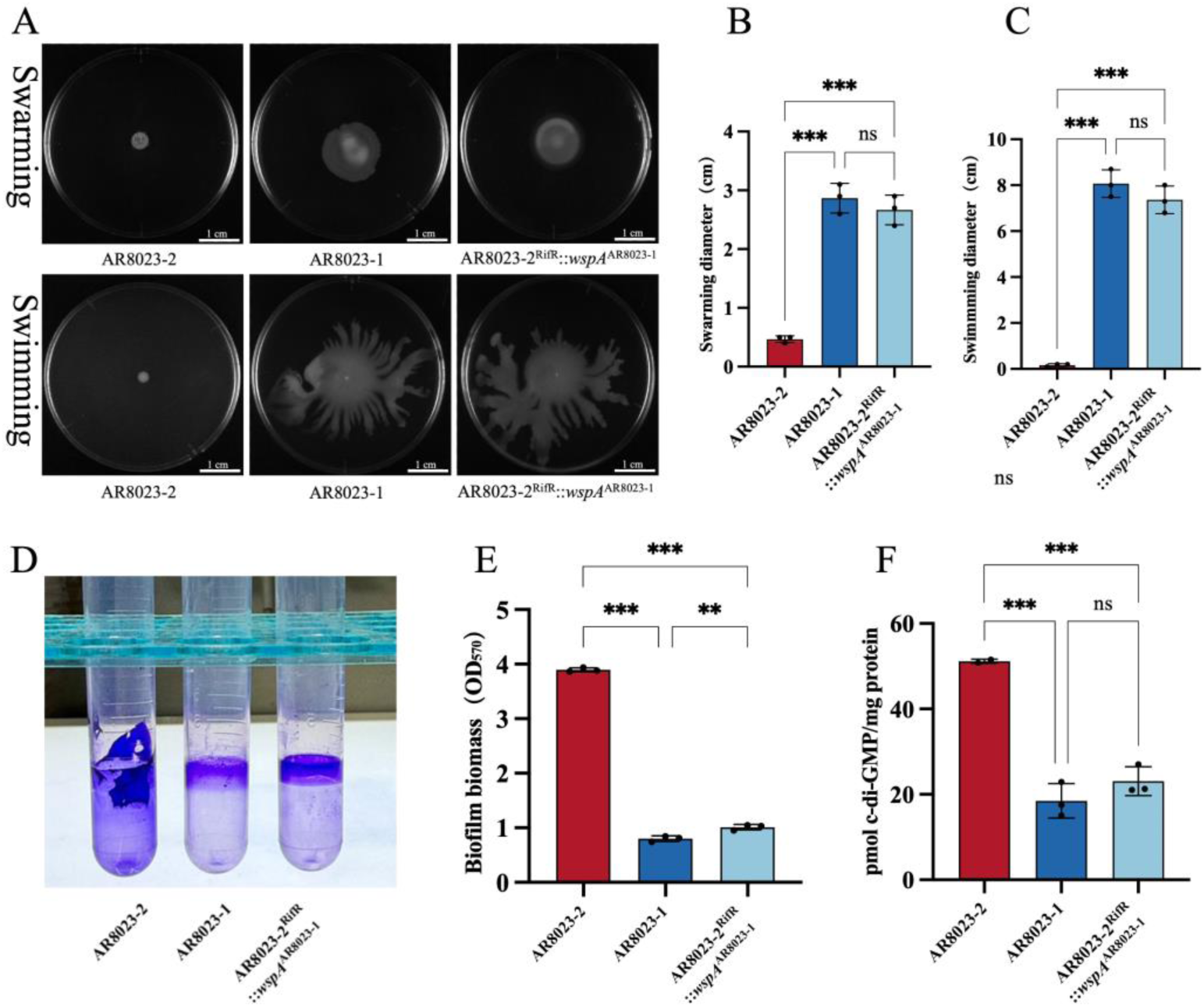
Assessment of motility phenotypes and determination of c-di-GMP and biofilm. (A) Representative images of swarming and swimming motility halos for the tested strains induced with 0% or 1% arabinose. (B) Quantification of swarming diameter (cm). (C) Quantification of swimming diameter (cm). (D) Morphology of the biofilm of the strain after staining with 1% crystal violet. (E) Biofilm productions of the strains grown in Luria-Bertani medium for 24 h. (F) The total intracellular c-di-GMP concentration of the tested strains was measured by LC/MS/MS after 24 h of growth. \*\**p* < 0.01, \*\*\**p* < 0.001, ns, non-significant.

Subsequently, to investigate the impact of bacterial biofilm on motility phenotypes, swarming and swimming motility were assessed using LB medium supplemented with gradient concentrations of agar (Fig. 2A). AR8023-2 exhibited significantly impaired swarming (0.47 ± 0.06 cm) and swimming (0.17 ± 0.06 cm) motility compared to AR8023-1 (2.87 ± 0.25 cm, *P* < 0.001; 8.07 ± 0.60 cm, *P* < 0.001) and AR8023-2^RifR^::*wspA*^AR8023-1^ (2.67 ± 0.25 cm, *P* < 0.001; 7.37 ± 0.60 cm, *P* < 0.001) (Fig. 2B-C).

### Transcriptomic profiling and c-di-GMP levels

To elucidate the mechanistic basis of phenotypic changes induced by the *wspA* mutation, transcriptomes were performed on clinical isolates AR8023-1 (control group A), AR8023-2 (experimental group), and the complemented strain AR8023-2^RifR^::*wspA*^AR8023-1^ (control group B) to identify differentially expressed genes (Fig. 3A-B). KEGG enrichment analysis of the differentially expressed genes revealed significant enrichment in pathways associated with biofilm formation. Notably, multiple up-regulated genes were implicated in the c-di-GMP signaling regulatory network (Fig. 3C). Consistent with prior evidence in *P. aeruginosa*, the chemoreceptor WspA within the Wsp system senses surface contact signals and activates the downstream diguanylate cyclase WspR, thereby stimulating c-di-GMP synthesis and inducing a planktonic-to-biofilm transition ^[8]^. The WspAΔ289Q mutation identified in this study likely disrupts WspA receptor function, leading to constitutive activation of the downstream c-di-GMP signaling pathway and ultimately driving the phenotypic alterations observed in the mutant strain AR8023-2.

**Fig. 3.**
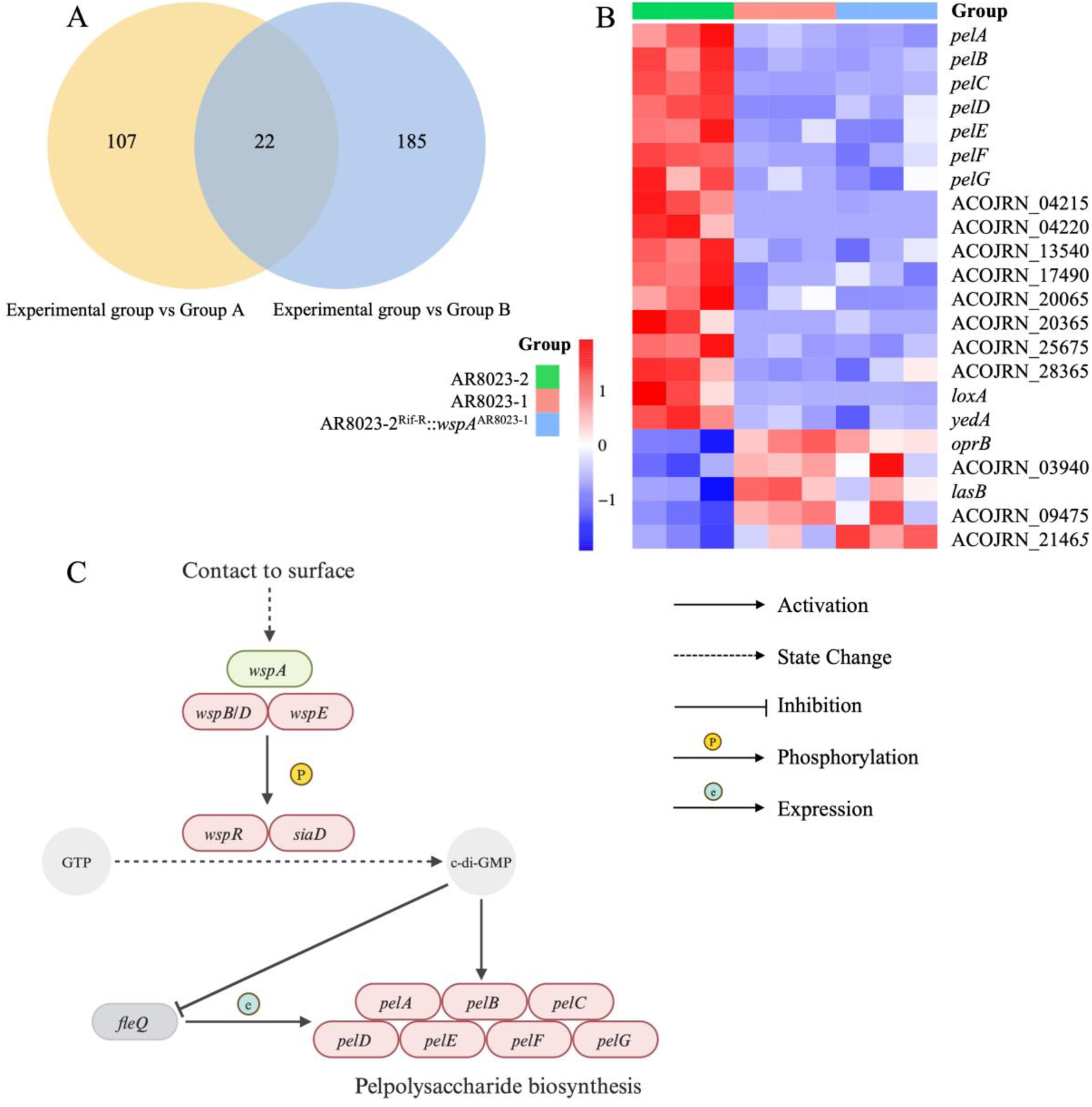
Transcriptomic profiling results. (A) The yellow circular region represents the differential gene set between the experimental group and the control group A, and the blue circular region represents the differential gene set between the experimental group and the control group B, where the intersection region (overlapping part) represents the differential gene set common to the two groups. (B) The expression patterns of the above 22 common differential gene sets were analyzed by heat map, and the color scale changes reflected the differential level of gene expression. (C) c-di-GMP signaling pathway, red is upregulated genes, and green is downregulated genes.

Transcriptomic profiling and prior literature indicate that the cyclic diguanylate monophosphate (c-di-GMP) signaling pathway is critically involved in biofilm formation. Therefore, HPLC-MS was employed to quantify intracellular c-di-GMP for all strains. The results demonstrated that c-di-GMP levels were ∼3-fold higher in AR8023-2 (51.15 ± 0.51 pmol/mg protein) versus AR8023-1 (18.50 ± 4.03 pmol/mg protein, *P* < 0.001) and AR8023-2^RifR^::*wspA*^AR8023-1^ (23.10 ± 3.39 pmol/mg protein, *P* < 0.001) (Fig. 2F).

### Enhanced antibiotic resistance

To systematically evaluate the impact of biofilm formation on antimicrobial susceptibility, the broth microdilution method (for planktonic strains) and MBEC Assay® (for biofilm strains) were employed. The susceptibility of the AR8023-1, AR8023-2, and AR8023-2^RifR^::*wspA*^AR8023-1^ to 10 antibiotics was assessed. The planktonic strains were designated as biofilm-low-producing strains for comparative analysis.

The results of the broth microdilution method demonstrated that all three strains (AR8023-1, AR8023-2, and AR8023-2^RifR^::*wspA*^AR8023-1^) remained susceptible to all tested antibiotics (Fig. 4A-B). In contrast, the results of the MBEC Assay® test revealed that the AR8023-2 exhibited Susceptibility to amikacin; Intermediate resistance to levofloxacin and ciprofloxacin; High-level resistance to ceftazidime, cefepime, meropenem, imipenem, ceftazidime-avibactam, and piperacillin-tazobactam, with MBIC values elevated ≥ 8- to 512-fold compared to planktonic MIC values (Fig. 4A-B). By comparing the antimicrobial susceptibility results of the two groups, we found that MBIC values of all strains exceeded planktonic MIC values by ≥ 4- to 64-fold except for amikacin and imipenem. Critically, the complemented strain (AR8023-2^RifR^::*wspA*^AR8023-1^) reverted to wild-type (AR8023-1) resistance levels in all assays, confirming that the WspAΔ289Q mutation directly correlates with altered biofilm-specific resistance phenotypes.

**Fig. 4.**
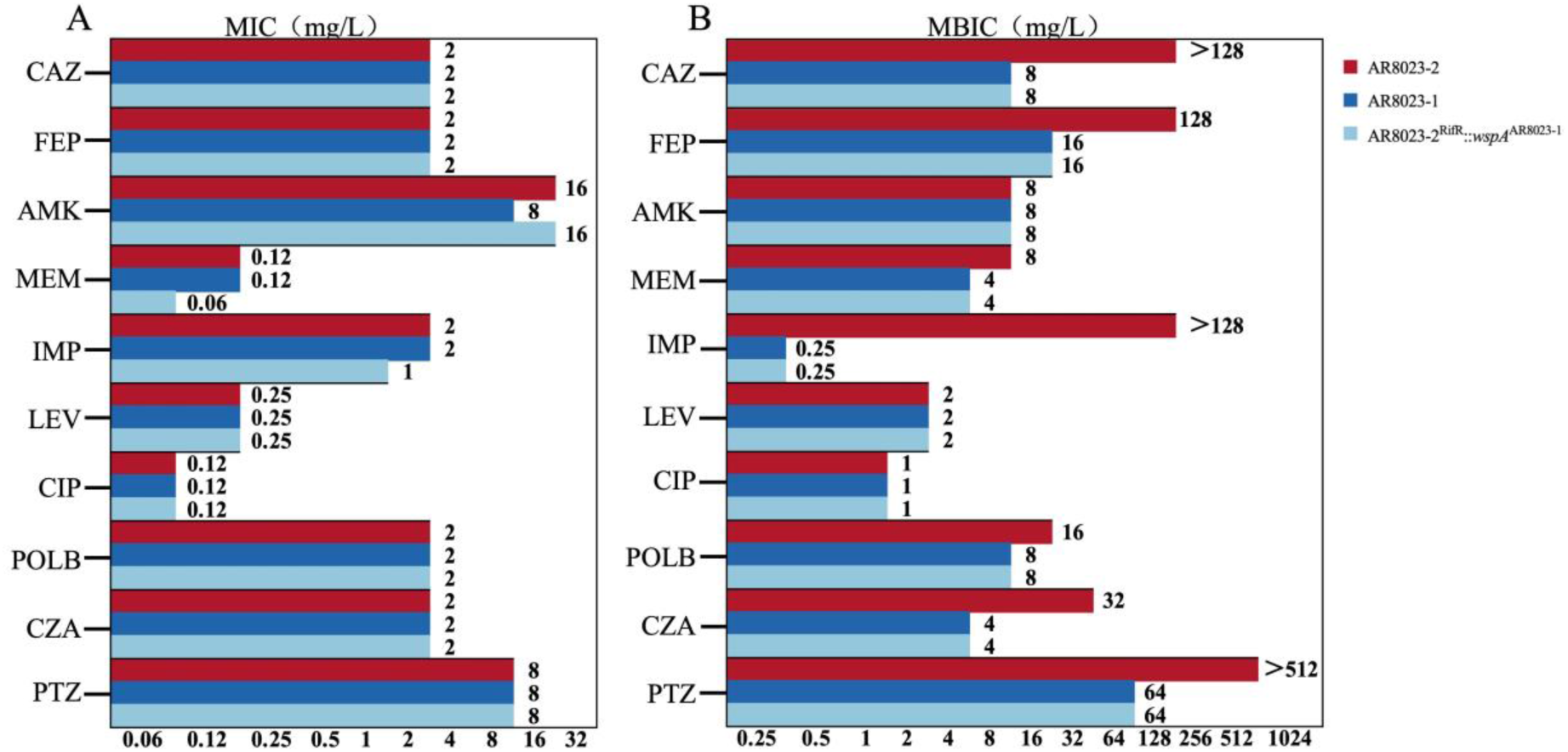
Results of antimicrobial susceptibility testing. (A) Planktonic MICs (for strains with low biofilm-forming capacity). (B) Biofilm MBICs (for strains with high biofilm-forming capacity).

## DISCUSSION

This study identifies a novel WspAΔ289Q mutation in clinical *P. aeruginosa* that drives RSCV formation through constitutive activation of the c-di-GMP pathway. The mutation disrupts WspA structural integrity, leading to aberrant phosphorylation of WspR and a near 3-fold increase in intracellular c-di-GMP levels (*P* < 0.001). Elevated c-di-GMP upregulates polysaccharide synthesis genes (e.g., *psl*, *pel*) ^[10,11]^, promoting hyper-biofilm formation and RSCV phenotypes while suppressing motility (*P* < 0.001). This discovery expands current understanding of WspA structural integrity in modulating colonial morphology and reveals a novel adaptive evolution mechanism in clinical *P. aeruginosa* isolates.

The Wsp chemosensory system in *P. aeruginosa*, a key network regulating surface attachment and biofilm formation ^[12–14]^, comprises six core components: the membrane-bound methyl-accepting chemotaxis protein WspA, membrane-anchoring scaffold protein WspB, signal transduction auxiliary proteins WspC and WspD, histidine kinase WspE, diguanylate cyclase WspR, and phosphatase WspF ^[7]^. As a transmembrane sensor, WspA activates the downstream diguanylate cyclase WspR through a phosphorylation cascade ^[5,15]^, thereby driving c-di-GMP synthesis ^[16,17]^. Previous studies identified a spontaneous in-frame deletion mutation (*wspA*Δ280-307) in the crude oil-isolated strain *P. aeruginosa* IMP68, resulting in a 28-amino acid deletion ^[8]^. This mutation removes two methylation sites (E280 and E297), preventing normal methylation modification of WspAΔ280-307. Consequently, this aberrant WspA variant constitutively activates WspR expression, leading to significantly elevated c-di-GMP levels. These findings underscore the critical role of structural integrity in WspA-mediated functional regulation.

Critically, the WspAΔ289Q mutation significantly enhanced biofilm-mediated antibiotic resistance. Biofilm MBICs for β-lactams (e.g., ceftazidime, meropenem) increased 8–512-fold compared to planktonic MICs, aligning with clinical treatment failure in the source patient. Importantly, RSCVs are intrinsically linked to persistent infections through enhanced biofilm formation and antibiotic tolerance. As a specialized form of small colony variants (SCVs), RSCVs exhibit metabolic dormancy and upregulated biofilm matrices that shield bacterial subpopulations from immune clearance and antimicrobial agents. This phenotype is conserved across pathogens like *Staphylococcus aureus* and *Pseudomonas aeruginosa*, where SCVs contribute to chronic infections by promoting antibiotic recalcitrance and relapse ^[18–20]^. Our identification of the WspAΔ289Q-driven RSCV pathway provides a molecular target for disrupting persistent biofilm infections, highlighting the translational relevance of c-di-GMP signaling inhibition in overcoming treatment failures[21].

Notably, this mutation’s phenotypic impact was strain-specific: introducing WspAΔ289Q into PAO1 did not induce RSCVs (Fig. S1), underscoring genetic heterogeneity in clinical isolates ^[22,23]^. Transcriptomics further revealed crosstalk between c-di-GMP and quorum sensing (QS), with dysregulation of *bapA* ^[24]^ (adhesion) and *lasA/lasB* ^[25]^ (matrix remodeling) contributing to biofilm resilience ^[26,27]^.

This study has the following limitations. First, the lack of an empty vector control strain (AR8023-2::pGK1900) complicates complementation validation. During its construction, this strain failed to grow on dual-antibiotic MHA plates (rifampicin 500 mg/L+gentamicin 50 mg/L), suggesting that the functional *wspA*-complementing plasmid may support essential metabolic functions absent in the empty vector. Second, model constraints limit physiological relevance: static in vitro cultures omit host factors (e.g., immune pressure, fluid shear stress); future studies should employ microfluidic ^[28]^ or animal infection models ^[29]^. Third, the single-isolate focus restricts generalizability; expanding screening to diverse clinical strains and correlating mutations with patient outcomes is essential. Finally, multi-omics depth requires enhancement: integrating metabolomics (e.g., TCA cycle remodeling ^[30]^) with spatial transcriptomics could resolve biofilm heterogeneity ^[31]^. Collectively, these limitations highlight needs for deeper exploration across model complexity, structural biology, and clinical translation.

The WspAΔ289Q mutation drives RSCV formation in clinical *P. aeruginosa* via constitutive activation of the Wsp-c-di-GMP pathway, leading to enhanced biofilm production, reduced motility, and increased biofilm-mediated antibiotic resistance (Fig. 5). This provides a molecular basis for RSCV persistence and informs strategies for targeted therapies and resistance surveillance.

**Fig. 5.**
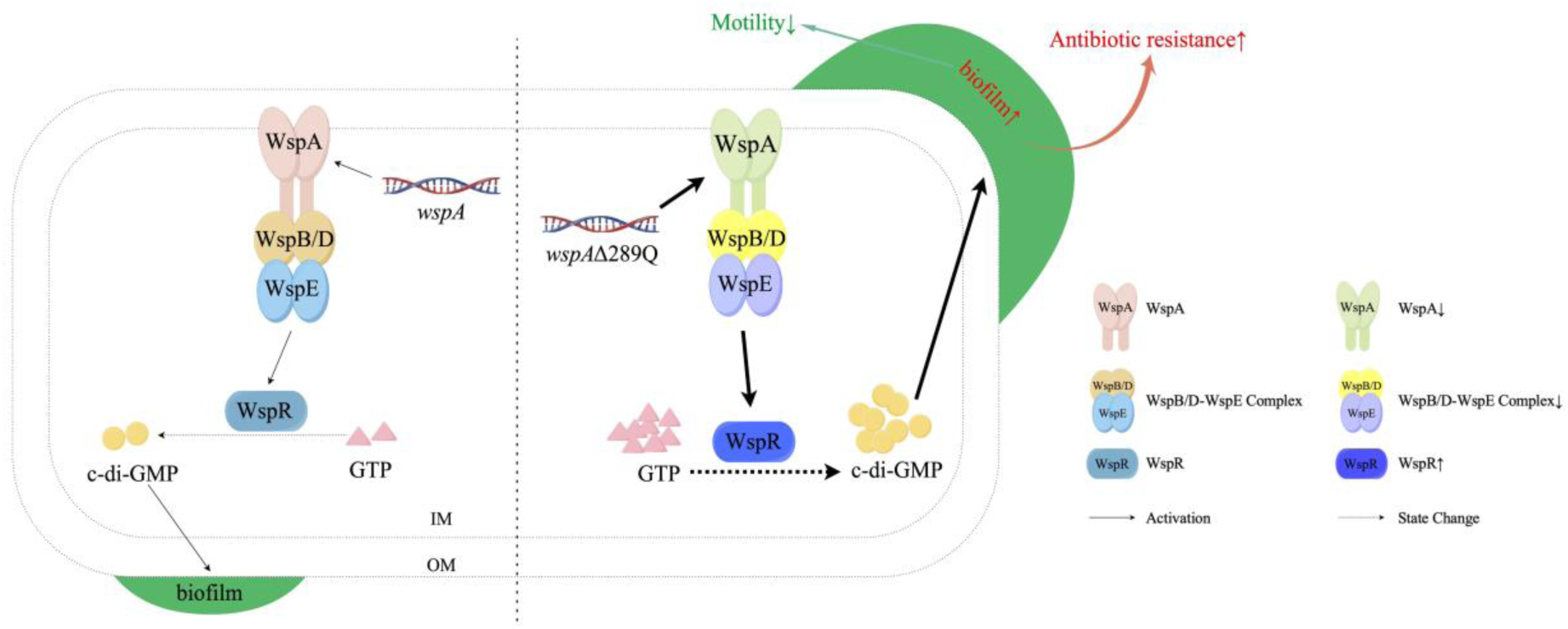
Proposed mechanism. Schematic of WspAΔ289Q mutation leading to constitutive WspR activation, elevated c-di-GMP, RSCV formation, enhanced biofilm, reduced motility, and increased antibiotic resistance.

## MATERIALS AND METHODS

### Bacterial strains and culture conditions

Clinical series strains and plasmids were listed in Table 1. PAO1 series strains and plasmids were listed in Table S1-S2. Unless specified, *Escherichia coli* and *P. aeruginosa* strains were routinely cultured in Luria-Bertani (LB) broth or Mueller-Hinton Agar (MHA) at 37°C. Antibiotics were supplemented at final concentrations as required: 300 mg/L carbenicillin, 100 mg/L tetracycline, 10 mg/L gentamicin, and 500 mg/L rifampicin for *P. aeruginosa*; 100 mg/L ampicillin, 10 mg/L gentamicin, and 50 mg/L kanamycin for *E. coli.* Colony morphology was assessed on modified LB agar plates (LBA; 10 g/L tryptone, 5 g/L yeast extract, 10 g/L NaCl, 15 g/L agar) supplemented with Congo Red (40 mg/L; sangon, China) and Brilliant Blue R (20 mg/L; sangon, China) to enhance visualization. This study was approved by ethics committee of Ningbo Medical Center Lihuili Hospital (approval number LY2024YJZ326).

**Table 1.** ***Pseudomonas aeruginosa* strains and plasmids used in the study.**

| Strains or plasmids | Relevant genotype and/or characteristics | Source |
| --- | --- | --- |
| *P. aeruginosa* |  |  |
| AR8023-1 | Clinical isolate (Wild-type) | This study |
| AR8023-2 | Clinical isolate (RSCV mutant) | This study |
| AR8023-2<sup>Rif<sup>R</sup></sup> | Rifampicin-resistant derivative of AR8023-2 | This study |
| AR8023-2<sup>Rif<sup>R</sup></sup>::*wspA*<sup>AR8023-1</sup> | AR8023-2<sup>Rif<sup>R</sup></sup> complemented with *wspA* from AR8023-1 via pGK_WspA<sup>8023-1</sup> | This study |
| Plasmids |  |  |
| pRK2013 | Helper plasmid for bacterial conjugation, Kan<sup>r</sup> | (Kyoung et al., 2006) |
| pGK1900 | *E. coli*–*Pseudomonas* shuttle vector, Gm<sup>r</sup> | (Li et al., 2021) |
| pGK_WspA<sup>8023-1</sup> | pGK1900 carrying *wspA* from AR8023-1, Gm<sup>r</sup> | This study |
| pGK_WspA<sup>8023-2</sup> | pGK1900 carrying *wspA* from AR8023-2, Gm<sup>r</sup> | This study |

Abbreviations: *P. aeruginosa*, *Pseudomonas aeruginosa*; *E. coli*, *Escherichia coli*; Gm, Gentamicin; Kan, kanamycin. The corner in the upper right corner of the gene indicates the strain from which the gene is derived.

### Strain construction

The WspA and its variants were cloned into pGK1900 plasmids as described previously ^[32]^, and the resulting expression vectors were subsequently introduced into *E. coli* DH5α. Then the recombinant plasmids were isolated, electroporation was used to introduce the recombinant plasmids into *P. aeruginosa* PAO1, and conjugation was used to introduce the recombinant plasmids into *P. aeruginosa* AR8023-2^RifR^ (rifampicin-resistant derivative of AR8023-2) ^[33]^.

The PAO1 series strains were constructed using CRISPR/Cas9-mediated genome editing ^[34]^. An in-frame deletion mutant (PAO1Δ*wspA*) was generated by cloning homologous arms flanking *wspA* into pACRISPR, transforming into PAO1 harboring pCasPA, and selecting mutants on tetracycline/carbenicillin plates, followed by plasmid curing via sucrose counter-selection. The site-specific ΔQ289 mutation was introduced into PAO1 via Gibson assembly of a repair template containing the CAG deletion and subsequent CRISPR/Cas9 editing, yielding PAO1*wspA*Δ289Q. For complementation, the WspAΔ289Q allele (with native promoter) from AR8023-2 was cloned into pGK1900 (Gm^R^), transformed into PAO1Δ*wspA*, and selected on gentamicin plates to generate PAO1Δ*wspA*::*wspA*Δ289Q^AR8023-2^. All mutants were verified by colony PCR and Sanger sequencing.

### Image acquisition of strains

The strains were inoculated on Colombian blood agar culture plates. After monoclones were grown, the plates were placed on a photography table, and photos were taken from front and side views, respectively.

Subsequently, the strains were streaked on MHA plates containing respective antibiotics to isolate single colonies (37℃, overnight). Single colonies were inoculated into LB broth with antibiotics and cultured at 37℃ with shaking (200 rpm, overnight). After ten-fold serial dilution to 10⁻⁶, 1 μL was spot-inoculated onto modified LB agar plates and incubated for 24 h at 37℃. Bacterial zones were imaged using a KEYENCE VHX-7000 stereomicroscope (KEYENCE, Japan) via triaxial adjustment, coarse focusing (20× objective), and fine focusing (200× objective) to capture high-resolution images.

### Whole genome sequencing and bioinformatics analysis

The genomic DNA of two clinical *P. aeruginosa* isolates was extracted using the QIAamp DNA Mini Kit (Qiagen, Germany) and subjected to Illumina short-read sequencing. De novo genome assembly was performed using shovill version 0.9.0 (https://github.com/tseemann/shovill). For further genomic analysis and experiments, we selected the AR8023-1 isolate for Nanopore sequencing. Hybrid assembly of long-read and short-read sequences was performed using Unicycler v0.4.8. Then, single nucleotide polymorphisms (SNPs) and insertion-deletion mutations were identified using Breseq (v 0.38.3) in standard analysis mode, with the complete genome of AR8023-1 serving as the reference sequence.

### Microtiter dish biofilm assay

Bacteria were grown on the MHA overnight and then inoculated by 100-times dilution into fresh LB broth in a 96-well plate (100 μl per well). After incubation at 37 ℃ for different times, biofilms formed on the inner wall of a well were stained with 0.1% crystal violet (CV) at 30 ℃ for 10 min. The dye was then dissolved with 30% acetic acid and spectrophotometrically measured at a wavelength of 570 nm.

### Motility assay

Swimming motility was assayed by stab-inoculating bacterial cells onto the surfaces of LB plates with 0.3% agar, followed by incubation at 37 ℃ overnight. Swarming assays were performed in LB plates with 0.5% agar and different concentrations of arabinose, as previously described ^[35]^. After inoculation, swarming plates were incubated for another 12– 24 h at 37 ℃ until the formation of swarming zones. Images were captured by Chemidoc XRS+, MAGELAB (USA).

### Transcriptome sequencing

*P. aeruginosa* AR8023-1, AR8023-2 and AR8023-2^RifR^::*wspA*^AR8023-1^were cultured overnight at 37° C in LB broth. Strains were diluted in 1:100 in fresh LB broth and grown at 37°C for 2 h. The 100 ml cells were collected at 4°C by centrifugation (5000 rpm,10 min). Total RNA was extracted using TRIZOL Reagent (USA) after liquid nitrogen grinding. Both wild type and mutants were prepared in bio-triplicate. Bacteria mRNA sequence library construction and sequencing were performed using Ribo-off rRNA Depletion Kit V2 (Vazyme, China). Raw data underwent stringent quality control (Trimmomatic), filtering adapter-contaminated reads (>5 bp), low-quality reads (Q < 15 in > 30% bases), and reads with >5% ambiguous bases (N), retaining high-quality data (Q20 > 97%, Q30 > 93%). Clean reads were aligned to the reference genome (BWA v0.7.13/Bowtie 2 v2.2.9), and gene expression was quantified as FPKM. Differential expression analysis (DESeq2) identified significant genes (|log₂FC|≥1, adj. *P* ≤ 0.05), with biological replicate consistency confirmed by Pearson correlation (> 0.85) and PCA. Functional enrichment of DEGs was conducted via BLAST against NR/KEGG/GO databases (E-value < 10⁻⁵), with pathway/term enrichment analyzed (topGO hypergeometric test, FDR≤0.05).

### Quantification of c-di-GMP

c-di-GMP (Sigma, SML1228, Germany) was extracted by heat/ethanol precipitation, quantified via HPLC-MS/MS (Agilent 6470, AB SCIEX QTRAP^®^6500; m/z 689→344 transition), and normalized to protein concentration by the following formula: N_c-di-GMP_=T_c-di-GMP_/T_protein_. ^[36]^. N_c-di-GMP_, Normalized c-di-GMP; T_c-di-GMP_, total c-di-GMP; T_protein_, total protein.

### Antimicrobial susceptibility testing

Antimicrobial susceptibility testing (AST) with ceftazidime (CAZ), cefepime (FEP), amikacin (AMK), meropenem (MEM), imipenem (IPM), levofloxacin (LEV), ciprofloxacin (CIP), polymyxin B (POLB), ceftazidime-avibactam (CZA), and piperacillin-tazobactam (PTZ) was performed using the broth microdilution method according to the Clinical and Laboratory Standards Institute (CLSI) guideline, and MIC breakpoints from the CLSI guideline M100 (2024) were used for the interpretation of susceptibility.

The MBEC (Minimum Biofilm Eradication Concentration) Assay^®^ kit (Innovotech, Canada) was used for antimicrobial susceptibility testing of biofilms. Due to the lack of uniform specifications and guidelines for the antimicrobial susceptibility testing of biofilm results, the minimal biofilm inhibitory concentration (MBIC) refers to the CLSI guideline M100 (2024). The antibiotics tested in this experiment are the same as the broth microdilution method.

### Statistical analysis

Statistical analysis and data visualization were performed using GraphPad Prism 10. Quantitative data are presented as mean ± standard deviation (SD). Differences between two groups were analyzed using Student’s t-test, while comparisons among multiple groups were assessed by one-way ANOVA followed by Tukey’s multiple comparisons test. A *P*-value of less than 0.05 was considered statistically significant.

## ACKNOWLEDGMENTS

This study was supported by the the National Key Research and Development Program of China 2023YFC2307100 (YY), the National Natural Science Foundation of China 82402665 (LZ), the Natural Science Foundation of Zhejiang Province LZY24H150003 (XH) and the Natural Science Foundation of Zhejiang Province LQN25H190002 (YL).

## AUTHOR CONTRIBUTIONS

Qing Yu, Conceptualization, Data curation, Formal analysis, Investigation, Methodology, Writing – original draft | Yue Li, Conceptualization, Funding acquisition, Investigation, Methodology, Writing – original draft | Peiqi Wang, Data curation, Investigation, Methodology | Chen Zhang, Methodology, Resources | Hanming Ni, Data curation | Yisha Zhang, Validation | Yushan Cui, Investigation | Mengyu Zhou, Conceptualization | Jun Ni, Data curation | Linyue Zhang, Funding acquisition | Xiaoting Hua, Funding acquisition, Writing – review and editing | Yunsong Yu, Project administration, Resources, Software, Supervision | Qiucheng Shi, Data curation, Methodology, Writing – review and editing | Xiaoxing Du, Project administration, Writing – review and editing |

## DATA AVAILABILITY

The sequence data supporting the findings of this study have been deposited in the National Center for Biotechnology Information under BioProject accession number PRJNA1308009.

---

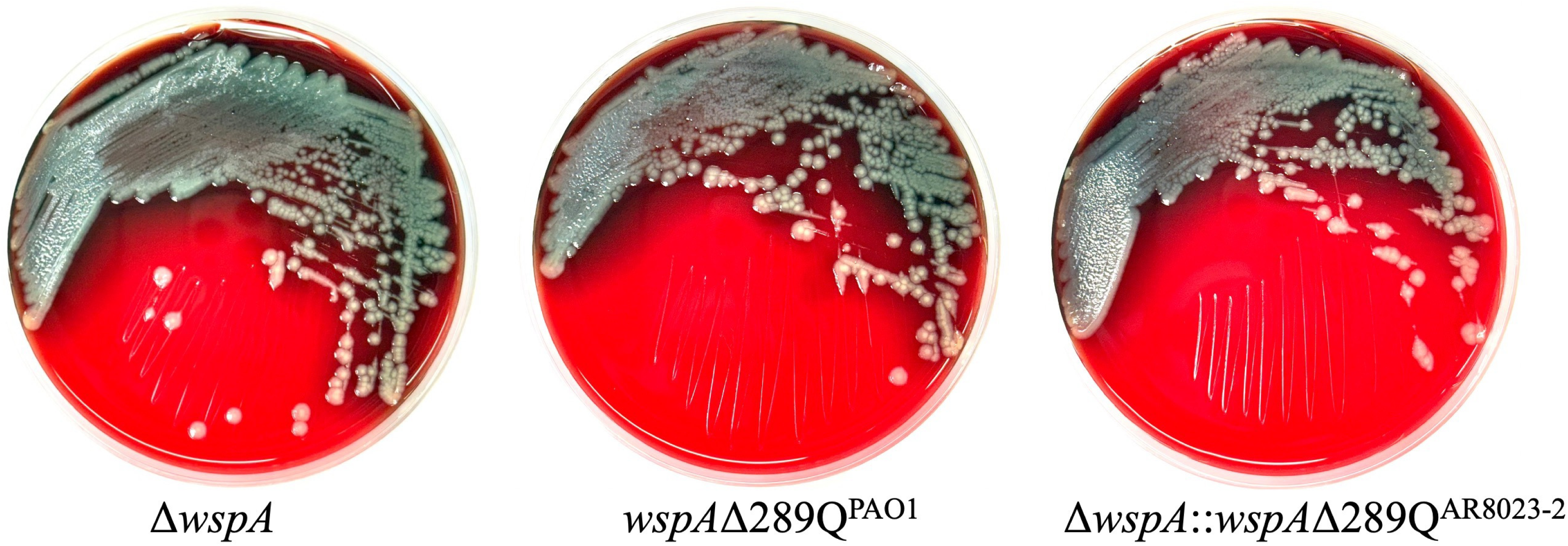

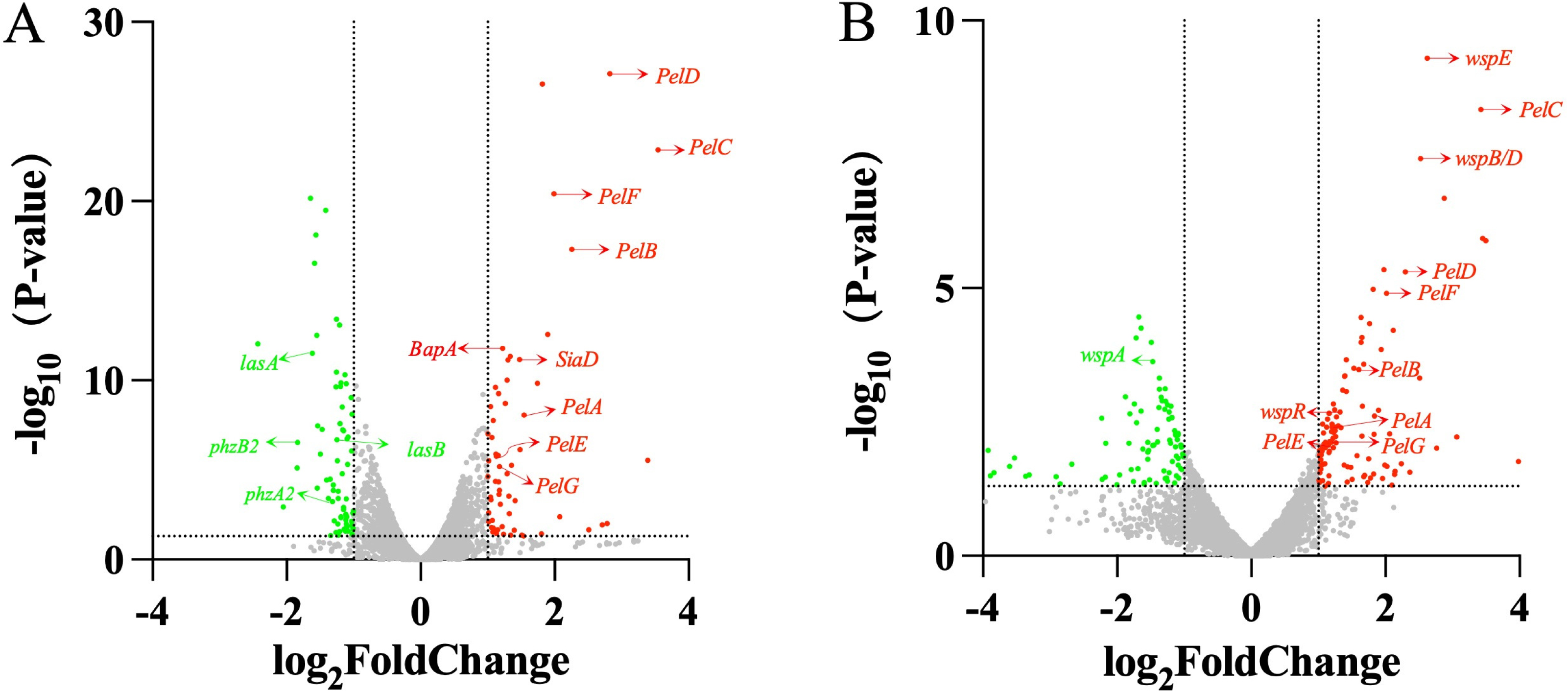

